# “Mine” or “yours”? Acoustic structure of food calls predicts sharing decisions in marmosets

**DOI:** 10.64898/2026.10.05.756650

**Authors:** Kaja Wierucka, Sven Winiger, Sandro Sehner, Judith M. Burkart

## Abstract

Foraging and feeding are often associated with vocalisations across animal species. Food calls may function to claim ownership of resources or recruit others to feeding sites, yet their potential role in facilitating food sharing remains poorly understood. This study investigates the relationship between food calls and food-sharing behaviour in common marmosets (*Callithrix jacchus*), focusing on adult–infant interactions, where food transfers are most frequent. We analysed over 1000 food interactions across six groups, categorising events as non-sharing (without begging or despite begging) or sharing (proactive, resisted, or facilitated). We examined the spectro-temporal structure of food calls produced immediately before each interaction. Food transfers were more likely to occur when preceded by a food call, suggesting a functional link between food calls and sharing. Random forest models reliably classified food calls according to interaction type, while generalized linear mixed models identified acoustic variation associated with sharing outcomes. Our results suggest that food calls encode information to either encourage or deter conspecific (infant) approach, influencing food transfer outcomes. These findings highlight the role of food calls in modulating social interactions around food in marmosets, advancing our understanding of vocal communication and its integration with cooperative behaviours in primates.

## Introduction

Vocalisations in the context of food acquisition and consumption are common across species. In primates, these food calls have been linked to a range of functions that vary across species and social contexts (Clay et al., 2012). One possible explanation for why food calls occur is that they simply reflect increased arousal in response to the possession of highly valuable food (Clay et al., 2012; Vitale et al., 2003). However, the absence of any social function is unlikely because calling is costly by making individuals susceptible to predation and must therefore be expected to be selected against. Indeed and consistent with the presence of a social function, food calls have been shown to be audience-dependent in several species (Vitale et al., 2003). In some species, such as white-faced capuchin monkeys (*Cebus capucinus*), food calls are emitted when others are in visual contact and the calls function to signal ownership of a food item (Gros-Louis, 2004). In other species, such as common marmosets (*Callithrix jacchus*), they are emitted when conspecifics are out of visual contact, attracting other individuals to the food source (Brown et al., 2004; Vitale et al., 2003). While this benefits conspecifics by informing them about the presence of food, it also reduces the caller’s own predation risk by attracting other individuals (Clay et al., 2012), thereby increasing group vigilance and lowering the caller’s probability of predation (Bednekoff & Lima, 1998; Lima & Dill, 1990). Some species, such as cooperatively breeding primates, emit food calls in the context of food sharing behaviour (Guerreiro Martins et al., 2019; Rogers et al., 2018; Troisi et al., 2018). Golden-lion tamarins, for instance, emit either a food-call that does not elicit an approach behaviour from conspecifics, or a food-offering call that encourages others to approach the caller (Boinski et al., 1994; Rapaport, 2006; Troisi et al., 2018). Whereas the former is either elicited due to higher arousal or in the context of providing information for individuals that have not yet detected the food (Boinski et al., 1994), the latter signals the willingness to share food with conspecifics (Rapaport, 2006; Troisi et al., 2018). Collectively, these patterns outlined above are consistent with the idea that non-human animals (henceforth animals) produce food-associated calls to regulate the social interactions around food.

Food sharing is commonly defined as a transfer of a defensible food item from one individual to another, with the exclusion of theft (Feistner & McGrew, 1989). It has evolved independently in several lineages of primates (Jaeggi & Van Schaik, 2011), and is underlain by different mechanisms in different species (Brown et al., 2004; Silk et al., 2013). Among nonhuman primates (henceforth primates), food sharing is most prevalent in cooperatively breeding callitrichid monkeys (tamarins and marmosets), occurring mainly, though not exclusively, between adults and immatures (Guerreiro Martins et al., 2019; Humle & Snowdon, 2008; Rapaport, 2006, 2011, 2020; Roush & Snowdon, 2001; Sehner et al., 2025; Troisi, 2021; Troisi et al., 2021). Callitrichid food sharing differs from that of other primates in two main respects. Firstly, food sharing in callitrichid monkeys is often unsolicited by the receiver and thus proactive from the side of the donor (Burkart & van Schaik, 2013; Guerreiro Martins et al., 2019; Sehner et al., 2025; Verspeek et al., 2022). This form of proactive sharing is most prevalent with immatures but also extends to unrelated and nonreciprocating individuals (Burkart et al., 2007) and even outgroup individuals (Troisi, 2021). Secondly, food-sharing in callitrichids may serve functions beyond mere nutritional provisioning (Brown et al., 2004). Several studies indicate that food sharing with naïve individuals (in particular from adults to immatures) serves the function to provide information about where and what to eat (Humle & Snowdon, 2008; Rapaport, 1999; Sehner et al., 2025; Troisi et al., 2018).

Common marmosets (*Callithrix jacchus*) are highly voluble callitrichid monkeys with a diverse vocal repertoire (Burkart et al., 2022; Schiel & Souto, 2017). Although their primary call types are fixed (as in all primates), marmosets exhibit considerable vocal flexibility by modifying call structure in response to their social environment. For example, they engage in vocal accommodation (Phaniraj et al., 2023, 2026; Zürcher et al., 2021; Takahashi et al. 2015, 2016, Wewhare et al. 2026) and exhibit population-level variation in call structure that amounts to vocal dialects (Zürcher & Burkart, 2017). Like other callitrichid monkeys, they often emit food calls when in the possession of a valuable food item (Guerreiro Martins et al., 2019; Vitale et al., 2003). Though food calls are produced more often when conspecifics are outside of visual contact (Vitale et al., 2003), adult marmosets also produce food calls in the presence of immatures, even when in visual contact (Brügger et al., 2018; Guerreiro Martins et al., 2019; Sehner et al., 2025). Especially in the first 20 weeks after birth, adult marmosets share a high proportion of their food with the immatures (Guerreiro Martins et al., 2019; Sehner et al., 2025), which is often accompanied by food calls. However, adult marmosets do not call for every food item they find, even very high-value ones, and a call does not guarantee that the item will be shared. This raises the question of whether food calls signal the willingness to share a food item. Food calls given in the presence of immatures may thus reflect heightened adult arousal that, via associative learning, draws immatures to approach and promotes sharing. Alternatively, they may also signal the propensity to share food. These mechanisms are not mutually exclusive and can operate alternately from one situation to another (i.e., an adult may be willing to share a first and a second highly valuable food item but decide to eat a third one themselves). Given the considerable acoustic diversity of marmoset food calls (Zürcher et al 2017., Agamaite et al. 2015), it may well be that these different goals are encoded in the acoustic features of the corresponding food calls.

In this study, we aimed to examine whether food call production and acoustic structure are systematically linked to food-sharing behaviour in common marmosets. First, we tested whether adults, after having acquired a highly valuable food item from a puzzle box, were more likely to share it with infants when producing food calls compared to when no call was uttered. We next examined whether the structure of food calls differed across sharing types, thereby testing whether the acoustic structure of food calls predict subsequent sharing decisions in marmosets. By addressing these questions, our study seeks to uncover the functional role of the broad acoustic variation in marmoset food calls and provide novel insights into how vocal communication is integrated with cooperative behaviours, advancing our understanding of the mechanisms by which marmosets regulate social interactions around food.

## Methods

### Subjects and housing

24 common marmosets across 6 different family groups were tested and recorded (14 adults and 10 infants; Supplementary materials, Table S1). All groups had infants at the time of the experiments. Animals were housed in temperature and humidity-controlled indoor enclosures (2.4 m × 3.2 m and 1.8 m) with access to an outdoor enclosure of the same size during appropriate outside temperatures. Both enclosures were filled with climbing structures, such as branches and ropes, and wooden chips and soil as floor texture. The animals received a diet of vitamin-supplemented rice flake mash in the morning, a vegetable mix at noon, and a protein-rich snack in the afternoon or gum arabica.

### Experimental design

The data used in this study were collected during experiments described by Sehner et al. (2022, 2023). Common marmoset family groups in their home enclosures were presented with three different food-extraction tasks. For each task, we presented a single apparatus containing eight crickets, with each cricket placed in a separate compartment and therefore requiring a separate extraction action. The action required was identical across all eight compartments within a given apparatus. In Task 1, individuals had to pull a sliding cover to access each cricket; in Task 2, they had to lift and hold up a box; and in Task 3, they had to remove a cotton plug from an Eppendorf tube and then invert the tube to retrieve the cricket (Supplementary Materials, Figure S1). A session was defined as a single presentation of one apparatus (5 minutes long), providing access to eight crickets. Each handling of a cricket was scored as one trial, beginning when an individual retrieved a cricket and ending upon complete consumption. Each task was presented to the animals in multiple sessions. Adult marmosets were habituated and trained on the tasks before the birth of an infant, while the infants were naïve to the experiment and therefore unable to solve these tasks (and did not learn to do so during the observation period). The experiments were recorded using a Handycam FDR-AX43 Camcorder, from which we extracted the audio data.

Only the first session of each testing day was included in the analyses to ensure comparable levels of food motivation among immatures, as earlier sessions often involved substantial food sharing and could therefore lead to varying degrees of satiation. In addition, only sessions containing at least one food-call bout were included. Session selection was independent of whether crickets were shared or not shared.

### Data processing

#### Video data

Videos were analysed using Mangold INTERACT software (Mangold, 2018). We recorded the identity of both the food retriever and the food consumer, the time stamp of each food call occurrence and the identity of the caller, as well as one of five food-interaction types (Figure 1). A cricket was classified as shared if it left the possession of the original food retriever and was directly obtained and consumed by another individual. Crickets that were dropped and later picked up by another animal were not considered shared. This definition, stricter than that proposed by Feistner & McGrew (1989), aligns with the highly social food-sharing behaviour of common marmosets (Guerreiro Martins et al., 2019).

**Fig. 1.**
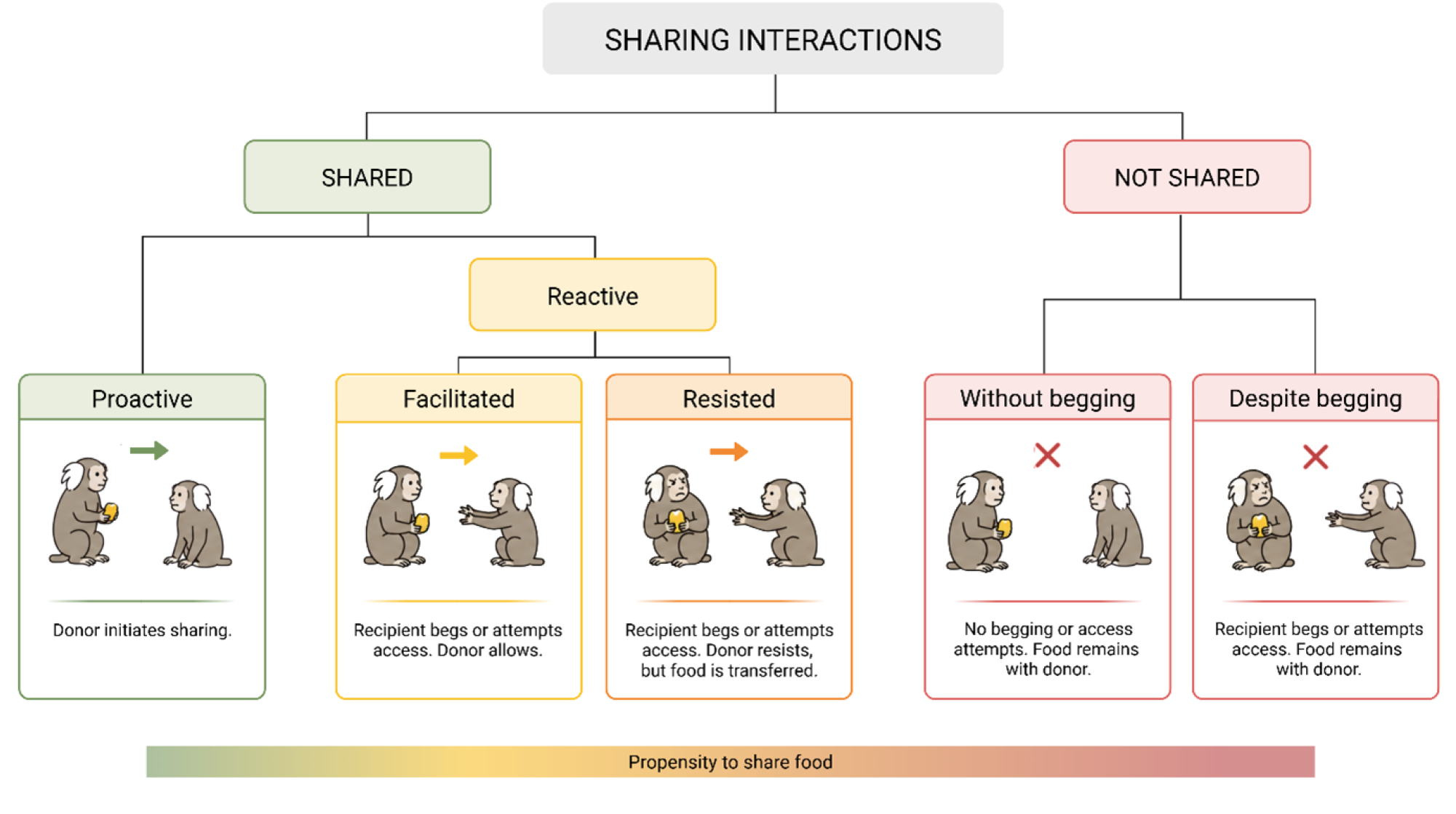
The different types of food interactions exhibited by common marmosets (following Sehner et al., 2023).

Food-sharing events were classified following Sehner et al. (2023). Sharing was first distinguished as proactive or reactive. In proactive sharing, the donor voluntarily initiated the transfer of the food item before the recipient approached or began begging, typically by presenting the food (e.g., holding it in an outstretched hand or mouth without consuming it) following the production of a food call. Reactive sharing was initiated by the recipient, defined as the recipient having approached, begged for, or attempted to access the food. Reactive sharing was further classified as facilitated, in which the donor tolerated or permitted the recipient’s attempts to obtain the food, or resisted, in which the donor actively attempted to retain possession of the food (e.g., by pulling it away, fleeing, or producing aversive vocalisations), but the food was nevertheless transferred. Non-sharing events were classified as occurring either without begging, when the recipient neither begged nor attempted to obtain the food and the original possessor retained it, or despite begging, when the recipient begged or attempted to access the food but the donor retained possession (Figure 1).

Interobserver reliability for the classification of the sharing behaviour was tested by comparing 10% of sessions coded by a second observer and a Cohen’s Kappa was calculated.

#### Acoustic data

We converted audio from the Handycam FDR-AX43 Camcorder into WAV files using the Shotcut software (Meltytech LLC) and used Raven Pro software (ver. 1.6.5) (Charif et al., 2010) to extract food calls from the recordings (as defined by Vitale et al. 2003, corresponding to sd-peeps in Agamaite et al. 2015). Only food calls produced by adults before the infants initiated approach were analysed, allowing the interaction to be interpreted from a sender’s perspective. This ensured that vocalisations preceded any behavioural response by the infant and could therefore be related to the subsequent outcome of the interaction.

Spectrograms were generated using a Hann window-type, size 400 samples, a Hop size of 200 samples, and an overlap of 50% (Figure 2). Using the time stamps, which were previously coded in Mangold INTERACT, the food calls were selected and matched to the caller and sharing behaviour. One call was defined as an utterance that was clearly visually separated from other utterances. Calls occurring within 0.3 seconds of each other were considered a food call bout (following Meshinska et al., 2024). Each food call bout was classified as “good,” “moderate,” or “bad” quality based on spectrogram clarity. A bout was categorized as good quality if all calls were clearly visible on the spectrogram, with no overlap from calls of other individuals or high-amplitude background noise affecting the fundamental frequency. Bouts were classified as moderate quality if at least one call’s fundamental frequency was obstructed by other calls or noise, but the total number of calls within the bout remained countable. If the number of calls per bout could not be determined due to obstruction, the bout was classified as bad quality.

**Fig. 2.**
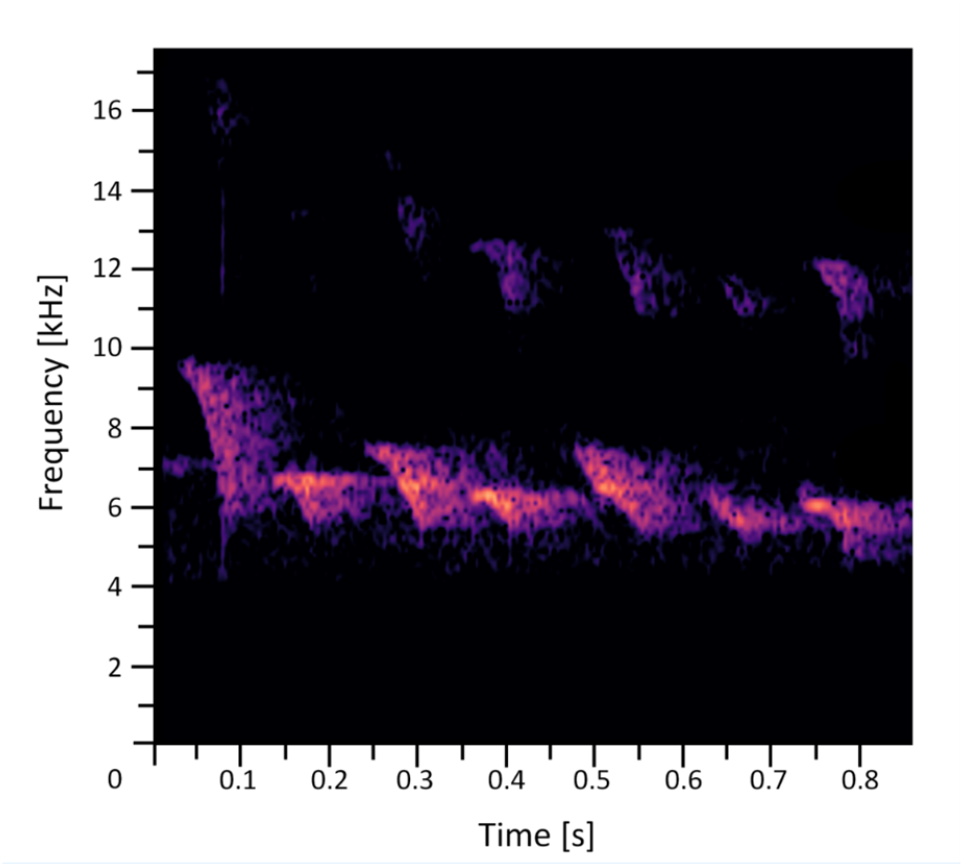
Spectrogram a food call bout. Spectrogram settings: Hann window-type, size 400 samples, a hop size of 200 samples, and an overlap of 50%. Bandstop filtered at 0-4000Hz.

To extract spectro-temporal features of the calls, we manually marked selection boxes around the peak harmonic on the spectrogram for each bout classified as moderate or good quality. Additionally, the lowest harmonic of each individual call within these bouts was selected. Bouts of bad quality and their respective calls were excluded from analysis. We then extracted robust acoustic measurements (Center Frequency, Frequency 5%, Frequency 95%, Frequency 25%, Frequency 75%, Bandwidth 50%, Duration 90%, Duration 50%) using Raven Pro (definitions in Charif et al., 2010). These features are based on the energy distribution within the call rather than the selection box itself, minimizing observer bias and enhancing measurement reliability.

### Statistical analysis

All statistical analyses were conducted in R ver. 4.3.2 (R Core Team, 2023). Generalized linear mixed models (GLMMs) were fitted using the lme4 package (Bates et al., 2024). To avoid overconfident parameter estimates and to maintain the nominal Type I error rate of 0.05, we implemented a maximal random-effects structure where supported by the data (Schielzeth & Forstmeier 2009; Barr et al. 2013). All variables were manually dummy coded and centered prior to inclusion as random slopes.

#### Sharing behaviour and food call production

We examined whether common marmosets share food more frequently with infants after producing a food call, using a binomial generalized linear mixed model (GLMM; lme4 package (Bates et al., 2024)). Whether a cricket was shared was included as a binomial response variable, and food call production (binomial) as a fixed effect, with individual identity nested in group as a random intercept.

#### Sharing behaviour and number of calls per bout

To test whether the number of calls per bout varied across food-sharing types, we fitted a GLMM with a negative binomial error distribution (to account for overdispersion). The number of calls per bout was included as the response variable, with food-sharing type and caller sex as fixed effects. Caller identity nested within group identity was included as random intercepts. We included random slopes for food-sharing type and sex within group identity, as well as random slopes for food-sharing type within caller identity nested within group identity. Bouts classified as poor quality were excluded from the analysis. Additionally, because callers may cease vocalizing abruptly when food sharing occurs, the final bout of each trial was excluded to avoid bias introduced by artificially shortened call bouts.

Multicollinearity was assessed using variance inflation factors (VIFs) calculated from a reduced model without interaction terms; the highest VIF was 1, indicating no issues with collinearity (Quinn & Keough 2002; Field 2005). Model stability was further assessed by sequentially removing individual levels of each random effect and comparing resulting estimates with those obtained from the full dataset (Nieuwenhuis et al. 2012; function provided by Roger Mundry). Fixed-effect estimates were considered acceptably stable. The model showed no evidence of overdispersion (dispersion parameter = 1.08).

To test the overall effect of food-sharing type on the number of calls produced per bout, we compared the full model with a reduced null model that excluded food-sharing type but retained all other terms.

#### Acoustic structure of calls and types of food interactions

We tested for overall differences in call acoustic structure across food-sharing types by assessing whether a random forest classifier could predict sharing type from the extracted acoustic features. We only included calls that had a sample size equal to or greater than 15 per individual per sharing type. To control for unequal sample sizes and potential individual-level biases, we balanced the dataset by randomly selecting 15 calls per individual per sharing type for analysis. Classification was performed using a modified version of the trainRF function from the GibbonR package (Clink et al. 2023). This modification split the data into non-overlapping training (70%) and testing (30%) sets while maintaining balance across individuals. Model performance was evaluated on the test set using confusion matrices generated with the confusionMatrix function from the caret package (Kuhn, 2008), from which classification accuracy was extracted. Because call selection was randomized and sample sizes were relatively small, the classification procedure was repeated 1,000 times to reduce stochastic bias. Reported accuracies represent the mean accuracy across all iterations. To determine whether classification performance exceeded chance levels, we adopted a one-sided statistical approach. Specifically, we calculated the 5th percentile of the accuracy distribution across iterations. Given five sharing types, chance-level accuracy was defined as 0.20. If the 5th percentile fell below this threshold, classification performance was considered not different from random. Conversely, if the entire distribution exceeded the random baseline, performance was deemed significantly above chance.

To further investigate how the acoustic structure of food calls varied with food-sharing type, we fitted linear mixed-effects models (LMMs). Prior to modelling, acoustic variables were reduced using principal component analysis (PCA), retaining components with eigenvalues greater than 1. Each retained principal component was then modelled separately using lmer from the lme4 package (Bates et al. 2024). Fixed effects included *food-sharing type* (five-level factor) and caller *sex* (two-level factor). In addition, based on visual inspection of calls and statistical comparisons between first and subsequent calls within bouts (Supplementary Figure S2), we included the relative position of the call within the bout (first vs. not first) and its interaction with food-sharing type. Random intercepts included call bout nested within caller identity nested within group identity. We included random slopes for food-sharing type and sex within group identity, as well as food-sharing type within caller identity nested within group identity. Model assumptions were evaluated visually using quantile–quantile plots and residuals-versus-fitted plots, with no clear violations detected. Model stability was assessed using the same procedures applied to previous models. Prior to model fitting, predictor and response distributions were examined; based on these inspections, PC1 and PC2 were log-transformed.

To test the effect of food-sharing type while avoiding issues of cryptic multiple testing (Forstmeier & Schielzeth 2011), we compared each full model to a reduced null model that excluded food-sharing type and its interaction with call order but retained all other fixed and random effects. Model-predicted values and 95% confidence intervals were estimated using parametric bootstrapping (1,000 iterations).

## Results

We processed data from a total of 1165 trials, from 18 individuals. The interobserver reliability for the coding of the sharing behaviour resulted in a Cohen’s Kappa of 0.72, which is considered a substantially strong agreement (Landis & Koch, 1977).

### Sharing behaviour and food call production

Following data cleaning (as described in the methods section) we used 1013 trials for analysis. Of the 1013 crickets that were handled, 288 were shared proactively, 211 facilitated, 46 resisted, 325 were not shared without begging, and 143 crickets were not shared despite begging, resulting in 545 shared crickets (53.8%) and 468 not shared crickets (Supplementary materials, Table S2). The adults (i.e. the potential donors) uttered food calls in 519 trials (51.2%). The binomial GLMM revealed that food sharing was more likely when food calls were produced (β=2.33, p < 0.001; Figure 3). When a food call was produced, the odds of food sharing were 10 times higher than when no call was produced.

**Fig. 3.**
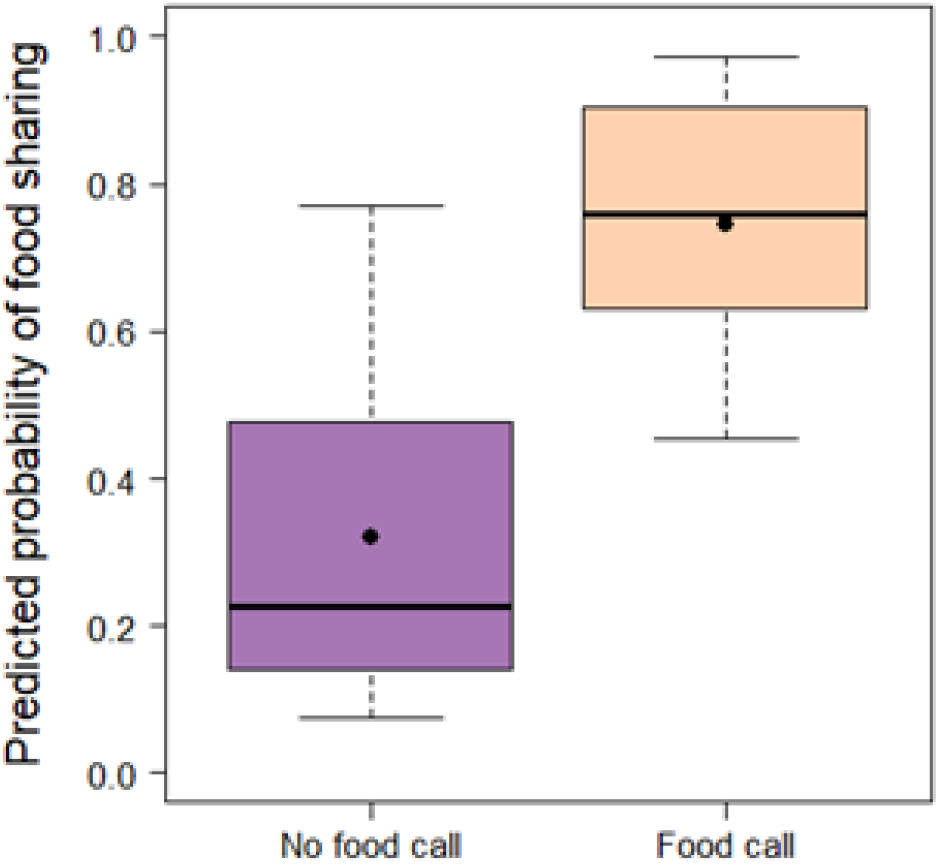
Effect of food calling on predicted probability of food sharing. Boxes (median and IQR) show predicted probabilities of food sharing (binomial GLMM) for trials with and without food calls. Black points indicate the mean predicted probability of food sharing for each condition.

### Sharing behaviour vs number of calls per bout

We analysed a total of 1121 bouts. The number of calls per bout ranged from 1 to 42. The GLMM unveiled that there was no effect of food sharing type on the number of calls per bout (2 = 2.136, p = 0.711).

### Acoustic structure of calls

We used 1888 calls for the acoustic structure analysis. After excluding instances with fewer than 15 calls per sharing type, 11 individuals remained in the analysis. The random forest model classified calls to food-sharing type with accuracies exceeding chance levels, with a mean accuracy of 0.358 (Figure 4).

**Fig. 4.**
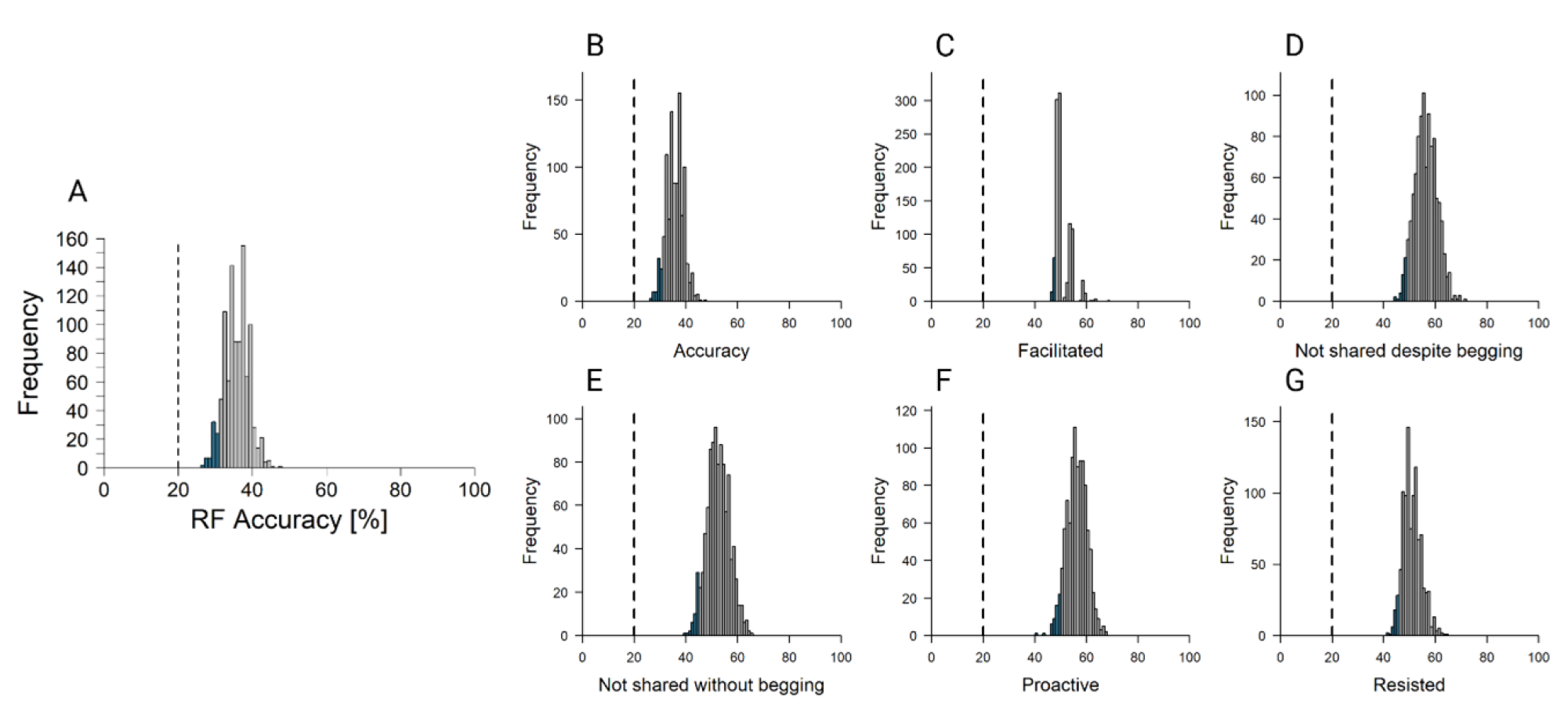
Distribution of random forest classification of marmoset food call acoustic features to food sharing type in which they were produced. The overall accuracy (A), as well as balanced accuracy for each class (food-sharing type; B-G) is shown, with the bottom 5th percentile in blue. The dashed vertical line represents the predefined random threshold. If the 5th percentile falls below this threshold, the model’s performance is not significantly different from random; otherwise, classification accuracy is considered better than chance.

The PCA returned 9 principal components, of which 3 had an eigenvalue greater than 1 (PC1: 4.811, PC2: 2.027, PC3: 1.812). Together, they explained 96.1% of the variation encountered in the original data, whereby principal component one (PC1) explained the highest amount with 53.5 %, followed by PC2 with 22.5% and PC3 with 20.1%. Examining the loadings of the PCA revealed that PC1 was mostly negatively correlated with frequency measurements, PC2 mostly positively with bandwidth measurement and negatively with Frequency 5%, and PC3 mostly negatively with duration measurements (Supplementary Materials, Table S3).

For PC1, comparison of the full and null models revealed a significant effect of food-sharing type and its interaction with call order on call acoustic structure (χ^2^ = 17.716, df = 8, p = 0.023). Because PC1 loaded primarily on frequency-related acoustic parameters, this indicates that frequency characteristics of food calls differed across food-sharing types. The interaction between food-sharing type and call order was marginally significant (F = 2.356, p = 0.052) and was therefore retained in the model.

First elements of bouts were higher-pitched than the rest of the elements in the bout (lower PC1 values, as frequency was negatively correlated; Figure 5). Facilitated sharing has the lowest frequencies (in both first and non-first elements), while when food was not shared (both without and despite begging), we observed the highest frequencies produced in non-first elements (Figure 5). The relative difference between first and non-first elements was largest for proactive as well as non-shared contexts (Figure 5).

**Fig. 5.**
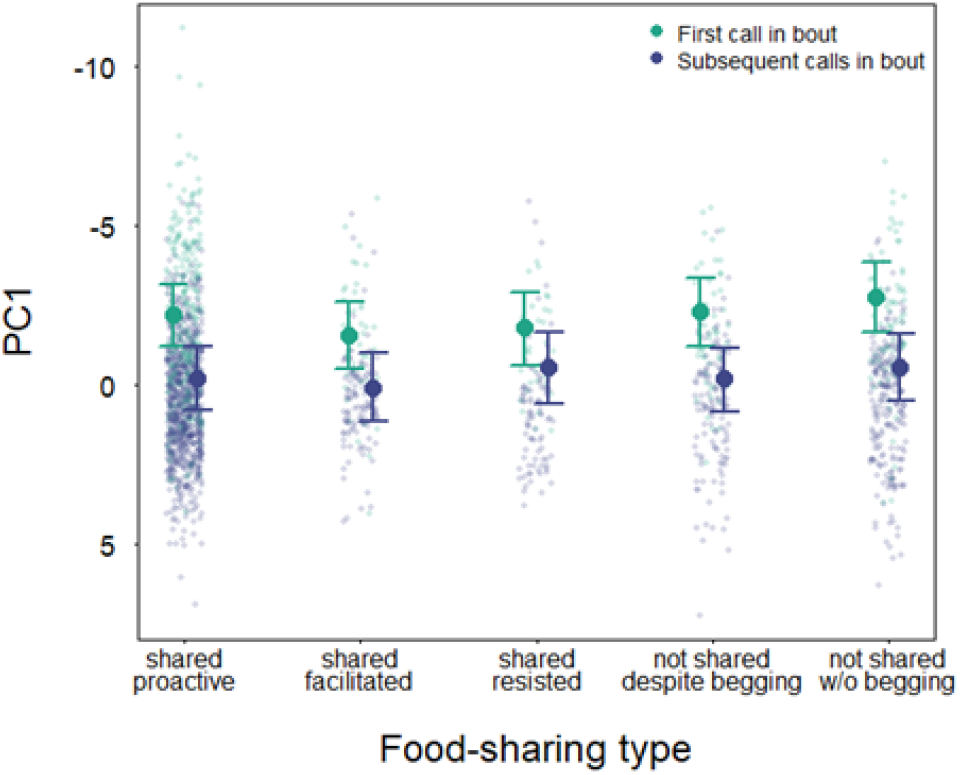
Differences in acoustic structure across food-sharing types in common marmosets. PC1 was negatively correlated primarily with frequency-related acoustic measures. Points show model-predicted mean PC1 values for each food-sharing type, separated by food-sharing order (first call of a bout in turquoise; subsequent calls in purple). Error bars indicate bootstrapped 95% confidence intervals. Grey points in the background represent raw PC1 values for individual calls.

In contrast, no effect of food-sharing type was detected for PC2 or PC3 (PC2: χ^2^ = 9.377, df = 8, p = 0.312; PC3: χ^2^ = 13.569, df = 8, p = 0.094), indicating that bandwidth- and duration-related features of call did not vary systematically across food-sharing types.

## Discussion

The function of food calls in common marmosets is largely unknown. In callitrichids more broadly, food-offering calls have been described as cues used to initiate food sharing with infants (Brown et al., 2004; Feistner & Price, 1990; Ferrari, 1987; Roush & Snowdon, 2001). However, common marmosets also produce food calls in contexts where food is not shared or is actively refused, suggesting that these calls may serve additional functions beyond simply advertising food availability. In this study, we investigated the relationship between food call production, their acoustic structure, and subsequent adult-to-infant food-sharing behaviour. Our results show that food calls are strongly associated with food-sharing events and that their acoustic structure varies systematically with the food-sharing type, implying that food calls may encode information about the caller’s propensity to share food.

We first examined whether overall, food call production was linked to food-sharing behaviour. Food sharing was substantially more likely to occur when a food call was produced compared to when no call was given, providing strong evidence for a link between food calling and food sharing in common marmosets. This relationship has long been assumed in the literature (Brown et al., 2004; Burkart & van Schaik, 2020; Guerreiro Martins et al., 2019; Sehner et al., 2023), but empirical evidence directly linking call production to distinct sharing outcomes was absent. Importantly, although food calls increased the likelihood of sharing, they were also produced in contexts where food was not shared. One possible explanation for this could be a lack of motivation or attention on the part of the infant, for instance due to satiation or distraction. However, food calls were also emitted in trials where the infant actively begged and the adult refused to transfer the food. These findings suggest that food calls cannot be interpreted solely as cues to invite approach, and instead raise the possibility that they convey more nuanced information related to sharing decisions.

The number of calls within a bout did not differ significantly across sharing types, indicating that call bout length did not influence whether or how food was shared. Previous work has shown that an increased number of calls per bout reflects heightened arousal in the caller (Borjon et al., 2016) and can increase arousal in a food-motivated receiver (Ermatinger et al., 2019; Owren & Rendall, 2011). The absence of an effect of bout size on sharing behaviour therefore suggests that neither the arousal state of the adult nor the arousal induced in the infant plays a significant role in determining sharing outcomes in this context. Instead, decisions about whether to share food appear to be governed by other mechanisms, potentially including the informational content of the calls themselves.

Analyses of call structure revealed clear differences between food calls produced in different sharing contexts. Random forest analyses indicated overall acoustic differentiation across sharing types when considering the full set of measured parameters. Generalized linear mixed models further showed that these differences were primarily driven by frequency-related features, while temporal features such as call duration and bandwidth did not vary with sharing behaviour. Specifically, food calls were lower-pitched in contexts where food was more likely to be shared, whereas higher-frequency calls were associated with non-sharing contexts. These findings suggest that frequency modulation may encode information about the caller’s willingness to share food. From a functional perspective, such acoustic variation could serve to either encourage or deter conspecific approach, thereby influencing the outcome of food-transfer interactions.

Importantly, the calls analysed here were produced by adults prior to infant approach behaviour, allowing the interaction to be interpreted from a sender’s perspective. The adult produced a food call, which was then available to the infant, who could decide whether or not to approach and beg. While it is not possible to determine whether infants always perceived or correctly decoded the acoustic differences in these calls, the behavioural outcomes are unambiguous: food was either shared or not, and these outcomes were associated with distinct acoustic patterns. If information about sharing propensity is valuable to the receiver, selection should favour signal variation that reflects the caller’s intention. The observed frequency differences between shared and non-shared contexts are consistent with this prediction. Notably, highest-frequency food calls were produced in trials where food was not shared and the infant did not beg, suggesting that infants may refrain from approaching because the adult had already encoded unwillingness to share in the call itself. Comparable functions have been described in other primates. Golden lion tamarins (*Leontopithecus rosalia*) produce two distinct food-related vocalisations. One is a food call that does not prompt conspecifics to approach an the other is a food-offering call that encourages others to join the food holder (Boinski et al., 1994; Rapaport 2006; Troisi et al., 2018). The food call potentially reflects arousal upon detecting food, whereas the food-offering call signals willingness to share. While common marmosets do not appear to produce distinct call types for this purpose, our findings suggest that they may achieve a similar outcome by modulating the frequency characteristics of a single call type. We thus find very nuanced communication about future food sharing decisions in two different species of cooperatively breeding primates, which is consistent with the idea that cooperative breeding facilitates cooperative communication among primates (Burkart et al. 2022).

We also found consistent acoustic differences between the first element of a call sequence and subsequent elements. This distinction is clearly audible and visible in spectrograms and may indicate that different elements within a bout serve different communicative roles. While the functional relevance of this pattern remains unclear, it highlights an additional layer of complexity in marmoset food calls and warrants further investigation, particularly with respect to how receivers attend to and process call sequences.

Several limitations of the present study should be acknowledged. First, our data were collected in a captive setting, which may constrain the generalisability of the findings to wild populations. However, previous work on wild marmosets suggests comparable patterns of food calling and sharing behaviour (e.g., de Oliviera Terceiro et al. 2021), indicating that the observed relationships are likely to extend beyond captive conditions. Second, although sex and breeding status differences are known to influence food-sharing dynamics in callitrichids, our dataset did not allow for a detailed examination of these factors, and future studies with larger samples should address how these differences interact with food call production and structure. Third, while we focused on variation in the acoustic structure of food calls in relation to sharing outcomes, additional sources of variation may arise from food type or quality. Experimental work by Rogers et al. (2018) has shown that marmosets can produce distinct food calls depending on the nature of the food (e.g., insects versus sweet fruits), raising the possibility that call structure encodes information about resource value. In the present study, this factor was controlled by using a single, highly preferred food item (crickets) presented in consistent quantities, thereby reducing confounding effects. Nonetheless, future research should explicitly investigate how variation in food type and nutritional value influences both food calling and the propensity to share, providing a more comprehensive understanding of the communicative and ecological drivers of these behaviours.

Taken together, these findings suggest that food calls in common marmosets seem to encode information about the propensity of the caller to share food. This points to a novel communicative function of food calls, whereby adults modulate vocal signals to manage infant approach and food transfer. These findings further support the notion that common marmosets flexibly adjust the acoustic structure of their food calls in ways that appear context-dependent, potentially reflecting mechanisms akin to vocal accommodation. Such flexibility has often been overlooked in non-human primates, yet growing evidence suggests it may play a more central role in the evolutionary foundations of human language than previously assumed (Stoll et al., 2025). In this light, the observed modulation of call frequency in relation to food-sharing outcomes may represent a form of graded signaling that allows individuals to influence social interactions with a high degree of nuance. Importantly, these results align with the hypothesis that cooperative breeding systems provide a socio-ecological context conducive to the evolution of more complex vocal coordination (Burkart et al., 2022). In species like marmosets, where infant provisioning and social tolerance are key, selection pressures may favour individuals capable of fine-tuning their vocal output to manage both affiliative and competitive aspects of feeding interactions. This supports the view that vocal communication in cooperative breeders is not only more frequent but also more functionally versatile, offering insights into how communicative complexity may emerge from the interplay between social structure and ecological demands.

## Data, code, and materials availability

The acoustic data and analysis code supporting the findings of this study will be made publicly available in an appropriate open repository upon acceptance of the manuscript for publication.

## References

Agamaite, J. A., Chang, C.-J., Osmanski, M. S., & Wang, X. (2015). A quantitative acoustic analysis of the vocal repertoire of the common marmoset (Callithrix jacchus). The Journal of the Acoustical Society of America, 138(5), 2906–2928. 10.1121/1.4934268

Barr, D. J., Levy, R., Scheepers, C., & Tily, H. J. (2013). Random effects structure for confirmatory hypothesis testing: Keep it maximal. Journal of Memory and Language, 68, 255–278.

Bates, D., Maechler, M., Bolker, B., & Walker, S. (2024). lme4: Linear mixed-effects models using Eigen and S4

Bednekoff, P. A., & Lima, S. L. (1998). Randomness, chaos and confusion in the study of antipredator vigilance. Trends in Ecology & Evolution, 13(7), 284–287. 10.1016/S0169-5347(98)01327-5

Boinski, S., Moraes, E., Kleiman, D. G., Dietz, J. M., & Baker, A. J. (1994). Intra-group vocal behaviour in wild golden lion tamarins, Leontopithecus rosalia : Honest communication of individual activity. Behaviour, 130(1–2), 53–75.

Borjon, J. I., Takahashi, D. Y., Cervantes, D. C., & Ghazanfar, A. A. (2016). Arousal dynamics drive vocal production in marmoset monkeys. Journal of Neurophysiology, 116, 753–764. 10.1152/jn.00136.2016

Brown, G. R., Almond, R. E. A., & van Bergen, Y. (2004). Begging, stealing and offering: Food transfer in non-human primates. Advances in the Study of Behaviour, 34, 265–295. 10.1016/S0065-3454(04)34007-6

Brown, K., & Mack, D. S. (1978). Food sharing among captive Leontopithecus rosalia. Folia Primatologica, 29(4), 268–290.

Brügger, R. K., Kappeler-Schmalzriedt, T., & Burkart, J. M. (2018). Reverse audience effects on helping in cooperatively breeding marmoset monkeys. Biology Letters, 14(3), 20180030. 10.1098/rsbl.2018.0030

Bugnyar, T., Kijne, M., & Kotrschal, K. (2001). Food calling in ravens: Are yells referential signals? Animal Behaviour, 61(5), 949–958. 10.1006/anbe.2000.1668

Burkart, J. M., Fehr, E., Efferson, C., & van Schaik, C. P. (2007). Other-regarding preferences in a non-human primate: Common marmosets provision food altruistically. Proceedings of the National Academy of Sciences, 104(50), 19762–19766. 10.1073/pnas.0710310104

Burkart, J. M., & van Schaik, C. P. (2013). Group service in macaques (Macaca fuscata), capuchins (Cebus apella) and marmosets (Callithrix jacchus): A comparative approach to identifying proactive prosocial motivations. Journal of Comparative Psychology, 127(2), 212–225. 10.1037/a0026392

Burkart, J. M., & van Schaik, C. P. (2020). Marmoset prosociality is intentional. Animal Cognition, 23(3), 581–594. 10.1007/s10071-020-01363-6

Burkart, J. M., Adriaense, J. E. C., Brügger, R. K., Miss, F. M., Wierucka, K., & van Schaik, C. P. (2022). A convergent interaction engine: Vocal communication among marmoset monkeys. Philosophical Transactions of the Royal Society B: Biological Sciences, 377(1859), 20210098. 10.1098/rstb.2021.0098

Chakrabarti, S., & Jhala, Y. V. (2017). Selfish partners: Resource partitioning in male coalitions of Asiatic lions. Behavioral Ecology, 28(6), 1532–1539. 10.1093/beheco/arx118

Charif, R., Waack, A., & Strickman, L. (2010). Raven Pro 1.4 user’s manual (11th ed.). Cornell Lab of Ornithology.

Clay, Z., & Zuberbühler, K. (2009). Food-associated calling sequences in bonobos. Animal Behaviour, 77(6), 1387–1396. 10.1016/j.anbehav.2009.02.016

Clay, Z., Smith, C. L., & Blumstein, D. T. (2012). Food-associated vocalizations in mammals and birds: What do these calls really mean? Animal Behaviour, 83(2), 323–330. 10.1016/j.anbehav.2011.12.008

Clink, D. (2023). gibbonR: An R package for the detection and classification of acoustic signals using machine learning (R package version 1.0.1).

de Kort, S. R., Emery, N. J., & Clayton, N. S. (2006). Food sharing in jackdaws, Corvus monedula : What, why and with whom? Animal Behaviour, 72(2), 297–304. 10.1016/j.anbehav.2005.10.016

de Oliveira Terceiro, F. E., Arruda, M. D. F., van Schaik, C. P., Araújo, A., & Burkart, J. M. (2021). Higher social tolerance in wild versus captive common marmosets: The role of interdependence. Scientific Reports, 11(1), 825. 10.1038/s41598-021-87672-5

de Waal, F. B. M. (1989). Food sharing and reciprocal obligations among chimpanzees. Journal of Human Evolution, 18, 433–459. 10.1016/0047-2484(89)90074-2

Di Bitetti, M. S. (2003). Food-associated calls of tufted capuchin monkeys (Cebus apella nigritus) are functionally referential signals. Behaviour, 140(5), 565–592.

Elowson, A. M., Tannenbaum, P. L., & Snowdon, C. T. (1991). Food-associated calls correlate with food preferences in cotton-top tamarins. Animal Behaviour, 42, 931–937. 10.1016/S0003-3472(05)80145-9

Ermatinger, F. A., Brügger, R. K., & Burkart, J. M. (2019). The use of infrared thermography to investigate emotions in common marmosets. Physiology & Behavior, 211, 112672. 10.1016/j.physbeh.2019.112672

Evans, C. S., & Evans, L. (1999). Chicken food calls are functionally referential. Animal Behaviour, 58(2), 307–319. 10.1006/anbe.1999.1143

Fedurek, P., & Slocombe, K. E. (2013). The social function of food-associated calls in male chimpanzees. American Journal of Primatology, 75(7), 726–739. 10.1002/ajp.22122

Feistner, A. T. C., & McGrew, W. C. (1989). Food-sharing in primates: A critical review. Perspectives in Primate Biology, 3, 21–36.

Feistner, A. T. C., & Price, E. C. (1990). Food-sharing in cotton-top tamarins (Saguinus oedipus). Folia Primatologica, 54, 34–45. 10.1159/000156424

Ferrari, S. F. (1987). Food transfer in a wild marmoset group. Folia Primatologica, 48, 203–206.

Field, A. (2005). Discovering statistics using SPSS. London: SAGE Publications.

Forstmeier, W., & Schielzeth, H. (2011). Cryptic multiple hypotheses testing in linear models: Overestimated effect sizes and the winner’s curse. Behavioral Ecology and Sociobiology, 65, 47–55.

Fox, J., & Weisberg, S. (2011). An R companion to applied regression (2nd ed.). London: SAGE Publications.

Goldstone, L. G., Sommer, V., Nurmi, N., Stephens, C., & Fruth, B. (2016). Food begging and sharing in wild bonobos (Pan paniscus): Assessing relationship quality? Primates, 57(3), 367–376. 10.1007/s10329-016-0522-6

Gros-Louis, J. (2004). The function of food-associated calls in white-faced capuchin monkeys, Cebus capucinus, from the perspective of the signaller. Animal Behaviour, 67(3), 431–440. 10.1016/j.anbehav.2003.04.009

Guerreiro Martins, E. M., Antonio, A. C., Finkenwirth, C., Griesser, M., & Burkart, J. M. (2019). Food sharing patterns in three species of callitrichid monkeys. Journal of Comparative Psychology. 10.1037/com0000169

Hamilton, W. D. (1964). The genetical evolution of social behaviour. I. Journal of Theoretical Biology, 7, 1–16. 10.1016/0022-5193(64)90039-6

Hauser, M. D. (1998). Functional referents and acoustic similarity: Field playback experiments with rhesus monkeys. Animal Behaviour, 55(6), 1647–1658. 10.1006/anbeh.1997.0712

Hauser, M. D., & Marler, P. (1993). Food-associated calls in rhesus macaques (Macaca mulatto): I. Socioecological factors. Behavioral Ecology, 4, 194–205.

Humle, T., & Snowdon, C. T. (2008). Socially biased learning in the acquisition of a complex foraging task in juvenile cottontop tamarins, Saguinus oedipus. Animal Behaviour, 75(1), 267–277. 10.1016/j.anbehav.2007.05.021

Jaeggi, A. V., & van Schaik, C. P. (2011). The evolution of food sharing in primates. Behavioral Ecology and Sociobiology, 65(11), 2125–2140. 10.1007/s00265-011-1221-3

Kitzmann, C. D., & Caine, N. G. (2009). Marmoset (Callithrix geoffroyi) food-associated calls are functionally referential. Ethology, 115(5), 439–448. 10.1111/j.1439-0310.2009.01622.x

Kuhn, M. (2008). Building predictive models in R using the caret package. Journal of Statistical Software, 28(5), 1–26. 10.18637/jss.v028.i05

Landis, J. R., & Koch, G. G. (1977). The measurement of observer agreement for categorical data. Biometrics, 33(1), 159–174.

Lima, S. L., & Dill, L. M. (1990). Behavioral decisions made under the risk of predation: A review and prospectus. Canadian Journal of Zoology, 68(4), 619–640. 10.1139/z90-092

Mangold, P. (2018). Discover the invisible through tool-supported scientific observation. In H. Böttger et al. (Eds.), Mindful evolution (pp. 1–16). Klinkhardt.

Meshinska, K., Burkart, J. M., Bell, M. B. V., & Wierucka, K. (2024). Echoes of self: Understanding acoustic structure and informational content in common marmoset phee sequences, BioRxiv: 10.1101/2024.04.14.589400

Michels, A. M. (1998). Sex differences in food acquisition and aggression in captive common marmosets. Primates, 39(4), 549–556. 10.1007/BF02557576

Nieuwenhuis, R., te Grotenhuis, M., & Pelzer, B. (2012). influence.me: Tools for detecting influential data in mixed effects models. The R Journal, 4, 38–47.

O’Bryan, L. R., Lambeth, S. P., Schapiro, S. J., & Wilson, M. L. (2021). Playbacks of food-associated calls attract chimpanzees towards known food patches. Primates, 62(6), 905–918. 10.1007/s10329-021-00936-4

Owren, M. J., & Rendall, D. (2011). Sound on the rebound: Bringing form and function back to the forefront. Evolutionary Anthropology, 10, 58–71.

Phaniraj, N., Wierucka, K., Zürcher, Y., & Burkart, J. M. (2023). Who is calling? Optimizing source identification from marmoset vocalizations with hierarchical machine learning classifiers. Journal of the Royal Society Interface, 20(207), 20230399. 10.1098/rsif.2023.0399

Phaniraj, N., Wierucka, K., & Burkart, J. M. (2026). Dynamic vocal learning in adult marmoset monkeys. Proceedings of the Royal Society B, 293(2026), 20260743. 10.1098/rspb.2026.0743

Quinn, G. P., & Keough, M. J. (2002). Experimental designs and data analysis for biologists. Cambridge University Press.

R Core Team. (2023). R: A language and environment for statistical computing. R Foundation for Statistical Computing. https://www.R-project.org/

Ramos, E. A., Kiszka, J. J., Pouey-Santalou, V., Ramírez Barragán, R., García Chávez, A. J., & Audley, K. (2021). Food sharing in rough-toothed dolphins. Marine Mammal Science, 37(1), 352–360. 10.1111/mms.12727

Rapaport, L. G. (1999). Provisioning of young in golden lion tamarins (Callithrichidae, Leontopithecus rosalia): A test of the information hypothesis. Ethology, 105(7), 619–636. 10.1046/j.1439-0310.1999.00449.x

Rapaport, L. G. (2006). Provisioning in wild golden lion tamarins (Leontopithecus rosalia): Benefits to omnivorous young. Behavioral Ecology, 17(2), 212–221. 10.1093/beheco/arj016

Rapaport, L. G. (2011). Progressive parenting behavior in wild golden lion tamarins. Behavioral Ecology, 22(4), 745–754. 10.1093/beheco/arr055

Rapaport, L. G. (2020). Social contributions to the foraging behavior of young wild golden lion tamarins (Leontopithecus rosalia): Age-related changes and partner preferences. American Journal of Primatology, 82(11). 10.1002/ajp.23056

Rogers, L., Stewart, L., & Kaplan, G. (2018). Food calls in common marmosets, Callithrix jacchus, and evidence that one is functionally referential. Animals, 8(7), 99. 10.3390/ani8070099

Roush, R. S., & Snowdon, C. T. (2001). Food transfer and development of feeding behavior and food-associated vocalizations in cotton-top tamarins. Ethology, 107(5), 415–429. 10.1046/j.1439-0310.2001.00670.x

Ruiz-Miranda, C. R., Kleiman, D. G., Dietz, J. M., Moraes, E., Grativol, A. D., Baker, A. J., & Beck, B. B. (1999). Food transfers in wild and reintroduced golden lion tamarins, Leontopithecus rosalia. American Journal of Primatology, 48(4), 305–320.

Ruiz-Miranda, C. R., Archer, C. A., & Kleiman, D. G. (2002). Acoustic differences between spontaneous and induced long calls of golden lion tamarins, Leontopithecus rosalia. Folia Primatologica, 73(2–3), 124–131.

Schiel, N., & Souto, A. (2017). The common marmoset: An overview of its natural history, ecology and behavior. Developmental Neurobiology, 77(3), 244–262. 10.1002/dneu.22458

Schielzeth, H. (2010). Simple means to improve the interpretability of regression coefficients. Methods in Ecology and Evolution, 1, 103–113.

Schielzeth, H., & Forstmeier, W. (2009). Conclusions beyond support. Behavioral Ecology, 20, 416–420.

Sehner, S., Willems, E. P., Vinicius, L., Migliano, A. B., van Schaik, C. P., & Burkart, J. M. (2022). Problem-solving in groups of common marmosets. PNAS Nexus, 1(4), pgac168. 10.1093/pnasnexus/pgac168

Sehner, S., Willems, E. P., van Schaik, C. P., & Burkart, J. M. (2023). The evolution of information donation. Research Square: 10.21203/rs.3.rs-2498407/v1

Sehner, S., Willems, E. P., Baumeyer, A., Davis, L., van Schaik, C. P., & Burkart, J. M. (2025). Sensitivity to immature skill deficits: Food sharing experiments in squirrel monkeys (Saimiri boliviensis) and common marmosets (Callithrix jacchus). Journal of Comparative Psychology, 139(3), 178–191. 10.1037/com0000399

Silk, J. B., Brosnan, S. F., Henrich, J., Lambeth, S. P., & Shapiro, S. (2013). Chimpanzees share food for many reasons: The role of kinship, reciprocity, social bonds and harassment on food transfers. Animal Behaviour, 85(5), 941–947. 10.1016/j.anbehav.2013.02.014

Slocombe, K. E., & Zuberbühler, K. (2006). Food-associated calls in chimpanzees. Animal Behaviour, 72(5), 989–999. 10.1016/j.anbehav.2006.01.030

Stevens, J. R. (2004). The selfish nature of generosity. Proceedings of the Royal Society B, 271(1538), 451–456. 10.1098/rspb.2003.2625

Stoll, S., van Schaik, C., Burkart, J. M., Ono, K., Phaniraj, N., Song, Z., & Bickel, B. (2025). From vocal homophily to vocal repertoire flexibility: Unravelling the socioecological drivers of language evolution. EcoEvoRxiv: 10.32942/X2V06N

Suzuki, T. N. (2012). Calling at a food source: Context-dependent variation in note composition of combinatorial calls in willow tits. Ornithological Science, 11(2), 103–107. 10.2326/osj.11.103

Takahashi, D. Y., Fenley, A. R., Teramoto, Y., Narayanan, D. Z., Borjon, J. I., Holmes, P., & Ghazanfar, A. A. (2015). The developmental dynamics of marmoset monkey vocal production. Science, 349(6249), 734–738.

Takahashi, D. Y., Fenley, A. R., & Ghazanfar, A. A. (2016). Early development of turn-taking with parents shapes vocal acoustics in infant marmoset monkeys. Philosophical Transactions of the Royal Society B: Biological Sciences, 371(1693), 20150370. 10.1098/rstb.2015.0370

Tardif, S. D., Smuncy, D. A., Abbott, D. H., Mansfield, K., Schultz-Darken, N., & Yamamoto, M. E. (2003). Reproduction in captive common marmosets. Comparative Medicine, 53(4), 364–368.

Troisi, C. A., Hoppitt, W. J. E., Ruiz-Miranda, C. R., & Laland, K. N. (2018). Food-offering calls in wild golden lion tamarins (Leontopithecus rosalia): Evidence for teaching behavior? International Journal of Primatology, 39(6), 1105–1123. 10.1007/s10764-018-0069-z

Troisi, C. A. (2021). Intergroup food transfers in wild golden lion tamarins (Leontopithecus rosalia). Primates, 62(1), 223–231. 10.1007/s10329-020-00846-x

Troisi, C. A., Hoppitt, W. J. E., Ruiz-Miranda, C. R., & Laland, K. N. (2021). The role of food transfers in wild golden lion tamarins (Leontopithecus rosalia): Support for the informational and nutritional hypothesis. Primates, 62(1), 207–221. 10.1007/s10329-020-00835-0

Vahed, K. (1998). The function of nuptial feeding in insects. Biological Reviews, 73(1), 43–78. 10.1017/S0006323197005112

Verspeek, J., van Leeuwen, E. J. C., Laméris, D. W., Staes, N., & Stevens, J. M. G. (2022). Adult bonobos show no prosociality. PeerJ, 10; e12849. 10.7717/peerj.12849

Vitale, A., Zanzoni, M., Queyras, A., & Chiarotti, F. (2003). Degree of social contact affects the emission of food calls in the common marmoset (Callithrix jacchus). American Journal of Primatology, 59(1), 21–28. 10.1002/ajp.10060

Watson, S. K., Townsend, S. W., Schel, A. M., Wilke, C., Wallace, E. K., Cheng, L., West, V., & Slocombe, K. E. (2015). Vocal learning in the functionally referential food grunts of chimpanzees. Current Biology, 25(4), 495–499. 10.1016/j.cub.2014.12.032

Wilkinson, G. S. (1984). Reciprocal food sharing in the vampire bat. Nature, 308(8), 181–184. 10.1038/308181a0

Wewhare, N., Burkart, J. M., & Wierucka, K. (2026). Sequence-level vocal convergence in common marmosets. bioRxiv. 10.64898/2026.03.20.713272

Zürcher, Y., & Burkart, J. M. (2017). Evidence for dialects in three captive populations of common marmosets (Callithrix jacchus). International Journal of Primatology, 38(4), 780–793. 10.1007/s10764-017-9979-4

Zürcher, Y., Willems, E. P., & Burkart, J. M. (2021). Trade-offs between vocal accommodation and individual recognisability in common marmoset vocalizations. Scientific Reports, 11(1), 15683. 10.1038/s41598-021-95101-8

