## Supplementary materials for "“Mine” or “yours”? Acoustic structure of food calls predicts sharing decisions in marmosets"

Table S1. Overview of study individuals. All breeders and helpers reached the age of adulthood. Guapa was first a helper in the Grappa group and later transferred to a new group, where she became a female breeder. Each group consisted of a breeding pair, at least one infant and potentially one or two helpers. One group did not contain a breeding male as he died shortly before the study started. Notations: M – male, F – female.

| Individual | Group | Sex | Status |
|---|---|---|---|
| Grappa | Grappa | F | breeder |
| Craken | Grappa | M | breeder |
| Guapa | Grappa/Guapa | F | helper/breeder |
| Ginger | Grappa | F | helper |
| Gina | Grappa | F | infant |
| Giotto | Grappa | M | infant |
| Ninja | Guapa | M | breeder |
| Guarana | Guapa | F | infant |
| Gimli | Guapa | M | infant |
| Jandira | Jandira | F | breeder |
| Wombat | Jandira | M | breeder |
| Jumbo | Jandira | M | infant |
| Nala | Nala | F | breeder |
| Narnia | Nala | F | helper |
| Nivana | Nala | F | helper |
| Nasa | Nala | F | infant |
| Nemo | Nala | M | infant |
| Nutella | Nutella | F | breeder |
| Jam | Nutella | M | breeder |
| Nocioletta | Nutella | F | infant |
| Nugatti | Nutella | F | infant |
| Wisconsin | Wisconsin | F | breeder |
| Tabor | Wisconsin | M | breeder |
| Wuschel | Wisconsin | M | infant |

Table S2. Number of analysed trials per sharing type per individual.

| Individual | shared facilitated | not shared despite begging | not shared without begging | shared proactive | shared resisted |
|---|---|---|---|---|---|
| Ginger | 6 | 8 | 9 | 2 | 2 |
| Grappa | 24 | 29 | 37 | 11 | 1 |
| Guapa | 3 | 2 | 28 | 5 | 0 |
| Jam | 17 | 7 | 6 | 21 | 0 |
| Jandira | 0 | 1 | 5 | 5 | 0 |
| Nala | 5 | 5 | 15 | 25 | 5 |
| Narnia | 1 | 6 | 5 | 3 | 1 |
| Ninja | 31 | 10 | 32 | 58 | 7 |
| Nirvana | 57 | 8 | 5 | 22 | 6 |
| Nutella | 43 | 10 | 38 | 51 | 16 |
| Tabor | 6 | 24 | 90 | 29 | 1 |
| Wisconsin | 4 | 7 | 10 | 18 | 0 |
| Wombat | 14 | 26 | 45 | 38 | 7 |
| **Total** | **211** | **143** | **325** | **288** | **46** |

Table S3. Loadings of the first three principal components. Highlighted are the loadings with an absolute value greater than 0.35.

| Parameter | PC1 | PC2 | PC3 |
|---|---|---|---|
| Center Frequency | **-0.447** | -0.093 | 0.022 |
| Frequency 5% | **-0.381** | -0.361 | -0.067 |
| Frequency 95% | **-0.446** | 0.085 | 0.092 |
| Frequency 25% | **-0.426** | -0.237 | -0.015 |
| Frequency 75% | **-0.453** | 0.022 | 0.059 |
| Bandwidth 90% | -0.177 | **0.588** | 0.222 |
| Bandwidth 50% | -0.155 | **0.605** | 0.184 |
| Duration 90% | -0.089 | 0.197 | **-0.672** |
| Duration 50% | -0.080 | 0.217 | **-0.669** |

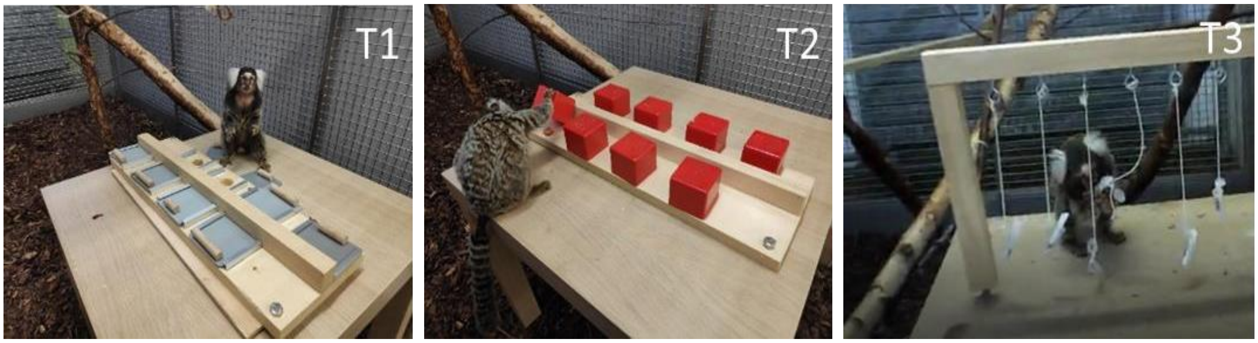


Figure S1. Experimental tasks presented to the common marmosets. (T1) Slider task: individuals had to pull a sliding cover to reveal the food reward beneath. (T2) Box task: individuals had to lift and hold up a box to gain access to the food reward. (T3) Tube task: food rewards were placed inside Eppendorf tubes sealed with cotton plugs. Individuals first had to remove the cotton plug and then invert the tube to retrieve the reward.

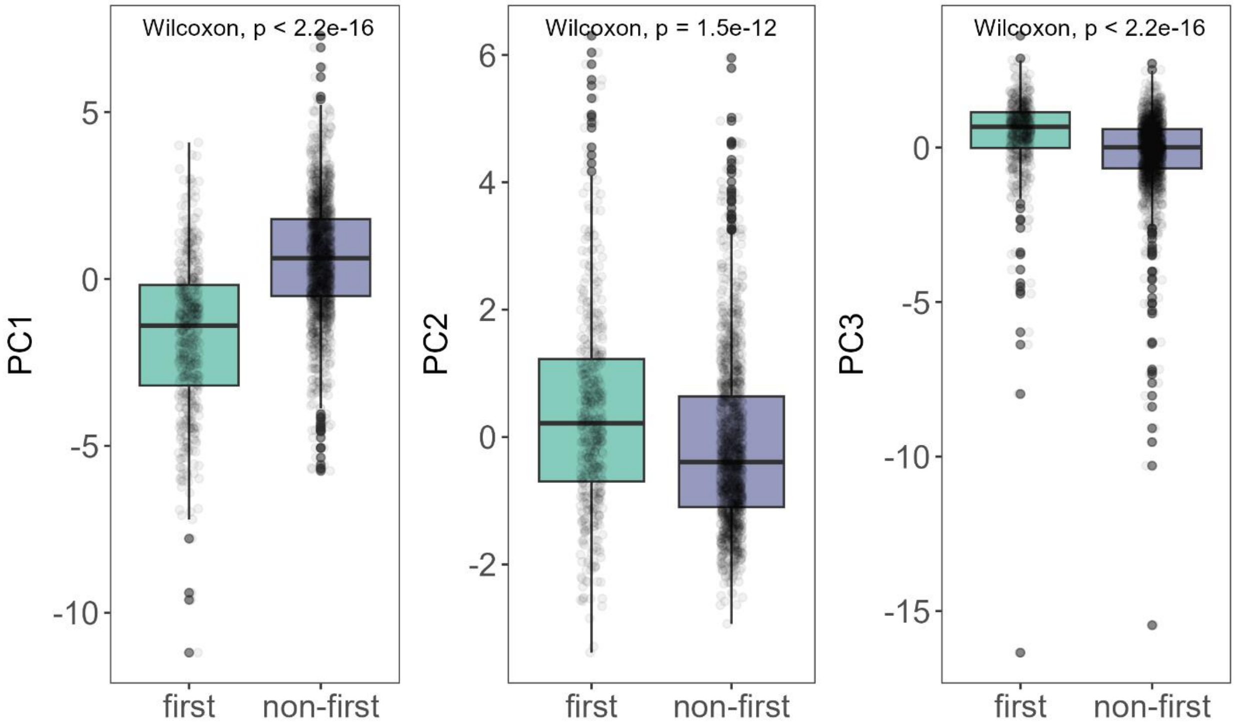


Figure S2. Structural differences between first and non-first calls in a bout. Principal component values (PC1, PC2, and PC3) are shown on the y-axis. Boxes represent the interquartile range, the horizontal line inside each box shows the median, and the whiskers extend to the most extreme points within 1.5 × IQR. Raw individual data points are plotted in the background. Statistical comparisons between first and non-first calls were performed using Wilcoxon rank-sum tests, with resulting p-values indicated above each boxplot.
